# Effect of exogenously applied alpha-tocopherol on vital agronomic, physiological and biochemical attributes of Lentil (*Lens culinaris* Medik.) under induced drought stress

**DOI:** 10.1101/2021.02.23.432440

**Authors:** Wadood Shah, Sami Ullah, Sajjad Ali, Muhammad Idrees, Muhammad Nauman Khan, Kasif Ali, Ajmal Khan, Muhammad Ali, Frhan Younas

## Abstract

Water being a vital part of cell protoplasm plays a significant role in sustaining life on earth; drastic changes in climatic condition leads to limit the availability of water and causing other environmental chaos. Alpha-tocopherol being a powerful antioxidant plays a vital role in scavenging the ill effects of oxidative stress. A pot experiment was conducted by exposing lentil cultivar (Punjab-2009) to varying levels of induced drought stress, sprinkled with α-tocopherol 100, 200 and 300 mg/L. Induced water deficit stress conditions caused a pronounced decline in growth parameters including absolute growth rate (AGR), leaf area index (LAI), leaf area ratio (LAR), root shoot ratio (RSR), relative growth rate (RGR), chlorophyll a, b, total chlorophyll content, carotenoids and soluble protein content (SPC) which were significantly enhanced by exogenously applied α-tocopherol. Moreover, a significant increase was reported in total proline content (TPC), soluble sugar content (SSC), glycine betaine (GB) content, endogenous tocopherol levels, ascorbate peroxidase (APX), catalase (CAT) peroxidase (POD) and superoxide dismutase (SOD) activities. On contrary, exogenously applied α-tocopherol significantly reduced the concentrations of malondialdehyde (MDA) and hydrogen peroxide (H_2_O_2_). In conclusion, it was confirmed that exogenously applied α-tocopherol under induced drought stress regimes ameliorated drought stress tolerance potential of lentil cultivar to a great extent; by enhancing growth, physiological and biochemical attributes.

## Introduction

Changing climatic condition is becoming an obstacle in fulfilling the demand of food and achieving a sustainable agriculture; climatic changes result in droughts, heavy floods, earthquakes, fluctuation in temperature and other environmental chaos that ultimately leads to reduce crop productivity [1]. Amongst the major abiotic stresses, drought stress has a profound effect on crop growth and yield reduction; though, plants can often withstand limited water condition but at the cost of substantial amount of plant total biomass and yield loss, drought stress condition disturbs vital physiological and biochemical processes ultimately declining plant growth and production [2]. About 50% of the world including arid and semi-arid being exposed to some kind of drought stresses [3]. With increasing climatic changes crops are losing their yield potential, thus making it hard to fulfil the increasing demand of food around the world [4]. The population of Pakistan is steadily increasing with 2.1% of the growth rate, which is greater than the growth rate (1.1%) of world population [5].

Abiotic stresses affect photosynthesis, cell growth, development and other vital physiological and biochemical processes [6]. According to previous research findings it is evident that water shortage in plants prompt oxidative stress, including both in the form of free radicals and non-radicals; both of the oxidants produced in response to abiotic stress damage biological membranes and other essential biomolecules such as proteins, lipids, chlorophyll and DNA [7]. Upon commencement of stress condition plants incline to accumulate a number of osmolytes such as; sugars, proteins, proline, glycine betaine, ornithine and citrulline that precisely play a role in osmoregulation: osmolytes are also known as compatible solutes having low-molecular weight accumulate chiefly in the cytoplasm preventing cellular degradation, because of their non-toxic nature and high solubility they do not impede other physio-bio chemical processes even if present in large amount [8].

In plants drought stress tolerance can be enhanced by using latest techniques like genetic engineering, plant breeding and tissue culture; however, these methods are comparatively expensive, time consuming and inflict adverse effects on health and environment; amongst different vitamins: α-tocopherol and nicotineamide are believed to be effectively alleviating the detrimental consequences of drought stress in plants by deactivating reactive oxygen specie (ROS) and shielding biological membranes against oxidative stress [9]. Encountering drought stress regimes plants trigger powerful antioxidant systems in the form of vitamins, flavonoids, carotenoids and antioxidant enzymes mainly including peroxidase, catalase, superoxide dismutase, glutathione reductase and ascorbate peroxidase [10].

Lentil (*Lens culinaris* Medik.) is an annual self-pollinated species belongs to family *Leguminacae* (*Fabaceae*) it is widely grown in South Asia, Middle East, North America, North Africa and Australia. Protein content of lentil seed ranges from 22% to 34.6% making it third highest level of protein of any legume or nut after soybeans and hemp [11].

The present research work was aimed to assess growth, physiological and biochemical responses of lentil cultivar (Punjab-2009) to varying levels of exogenously applied α-tocopherol, its potential in scavenging the negative impacts of induced drought stress and to explore the degree of efficacy of α-tocopherol in regulating vital metabolic processes by ameliorating drought tolerance potential of lentil cultivar subjected to varying levels of induced drought stress.

## Materials and Methods

### Site description and experimental design

Field experiment was carried out at the department of botany, University of Peshawar (34° 1’ 33.3012’’ N and 71° 33’ 36.4860’’ E.) Pakistan, during the growing season 2019. Peshawar is situated in Iranian plateau area having tropical climate. Soil texture was determined as sandy loam; evaluated via hydrometer method by Gee and Bauder, [12].

The seeds of lentil (*Lens culinaris* Medik.) variety punjab-2009 were obtained from National Agriculture Research Centre (NARC) Islamabad, Pakistan. Surface sterilized seeds were sown 2-4 centimetres deep in the soil filled earthen pots (20cm height, 18cm upper/lower diameter and 2cm thickness) contained 3kg sand and silt with a ratio of 2:1. Experiment was conducted in Randomized Complete Block Design (RCBD). Thinning and weed eradication were properly maintained and seedlings were exposed to sunlight for better growth. Three replicates were taken for each group. All the groups were normally watered till 60^th^ day of emergence. Different levels (100, 200 and 300 mg/L) of α-tocopherol were prepared by mixing 100, 200 and 300 mg tocopherol separately in 900 ml distilled water and 70% ethanol followed by heating at 33 °C for 15 minutes. After 60 days of germination one set of trial was exposed to 20-days of induced drought stress and sprayed with three levels of α-tocopherol (100, 200, and 300 mg/L) once throughout the growing season. Whereas, the second set of experiment was subjected to 25-days of induced drought stress and sprayed with the same levels of exogenously applied α-tocopherol. At the end of induced drought stress period’s five plants from each replicate were harvested randomly for the determination of various growth and physio-biochemical parameters.

### Soil analysis

Over the last two years average soil chemical properties were; Electrical conductivity (EC) 2.67 ds/m, pH 6.3 [13], Nitrogen (N) content 4.02g/kg [14], organic Carbon (C) 23.3 g/kg [15], potassium (K) available 91.4 mg/kg [16] and Phosphorus (P) 8.1 mg/kg [17].

### Growth measurements

Growth parameters; absolute growth rate (AGR), relative growth rate (RGR), coefficient of velocity of germination (CVG) and net assimilation rate (NAR) were calculated by following the formulas suggested by Ghule et al. [18].

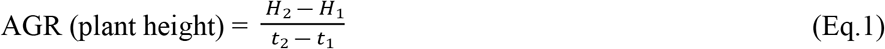

H_1_ and H_2_ denoted plant height (cm) during the time T_1_ to T_2_.

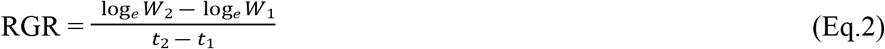

W_1_ and W_2_ denoted plant dry weight (gm) at time t_1_ and t_2_. Log^e^ is natural logarithm

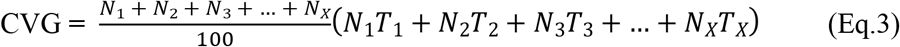

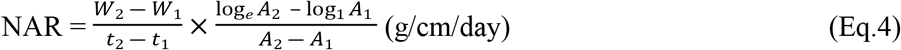

A_1_ and A_2_ denoted surface area of leaf and W_1_ and W_2_ are plant total dry matter at Time T_1_ and T_2._

Crop growth rate (CGR), leaf area index (LAI), leaf area ratio (LAR) and relative water content (RWC) were calculated by the formula suggested by Shah et al. [19].

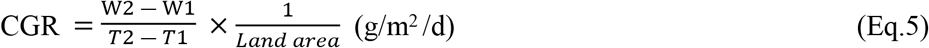

W_1_ and W_2_ are plant dry weights taken at time T_1_ and T_2_, respectively.

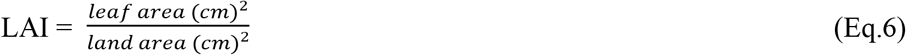

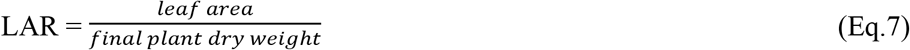

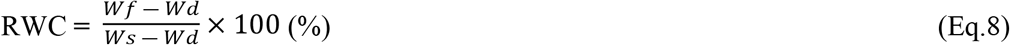

“Wf” represented leaf fresh weight and “Wd” leaf dry weight. “Ws” indicated saturated weight of leaf material determined after floating the leaves in distilled water for 18 hours.

Root-shoot ratio (RSR) was calculated by applying the formula proposed by Chuyong and Acidri, [20].

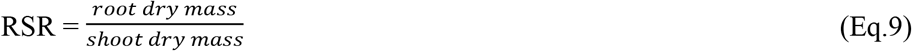

Seed vigor index (SVI) was measured by the formula proposed by Bina and Bostani, [21].

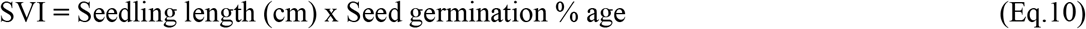

### Photosynthetic pigments (Chlorophyll a, b & carotenoids)

Fresh leaf material (0.5 gm) was homogenized in 80% acetone solution (10 ml) followed by centrifugation for 5 minutes. The samples were kept in dark overnight at 4°C. Finally, Optical density was measured at 470, 645 and 663 nm for carotenoids and chlorophyll a & b contents by following the protocol of Arnon, [22].

### Soluble sugar content (SSC)

Foliar material (0.5 gm) was grounded in 5 ml distilled water. The homogenized mixture was centrifuged for 10 minutes; after centrifugation 35% concentrated H_2_SO_4_ (4 ml) was added to 1ml supernatant. Optical density (OD) was noted at 490 nm by the methodology of Dubois et al. [23].

### Total proline content (TPC)

Proline content was quantified by the methodology of Bates et al. [24]. Fresh leaves (0.5 gm) were grounded in 10 ml 3% aqueous sulphosalicylic acid. The mixture was filtered and 2 ml filtrate was taken; 4 ml ninhydrin solution and 4 ml glacial acetic acid (20%) were mixed with filtrate. The mixture was heated at 100 °C for 1 hour and 4 ml toluene was added to it. OD readings were recorded at 520 nm.

### Glycine betaine content (GBC)

Fresh foliar material (0.5gm) was chopped in 10 ml distilled water. The mixture was filtered; filtrate obtained was diluted by adding 2 ml H_2_SO_4_ solution. The samples were centrifuged for 10 minutes and Cold KI–I_2_ was added to supernatant. 1 ml supernatant was collected and optical density was measured at 365 nm by using the methodology of Di-martino et al. [25].

### Soluble protein content (SPC)

Protein content in leaf tissues were investigated by following the protocol of [26]. 0.5 gm fresh leaf tissues were grounded in 5 ml phosphate buffer (pH 7.0) in ice cooled pestle and mortar. After grinding, the mixture was centrifuged for 15 minutes. 0.1 ml supernatant was taken and 2 ml Bradford reagent was added to it. Optical density was measured at 595 nm.

### Malondialdehyde content (MDAC)

Fresh leaf material (0.25 gm) was chopped in 3 ml 1.0% (w/v) Trichloro acetic acid (TCA). The mixture was spun in centrifuge machine for 10 minutes. 1 ml supernatant was taken and 4 ml 0.5% (w/v) 2-thiobarbituric acid was added. Samples were heated at 95 °C for 1 hour and then cooled by placing in ice bath for 10 minutes. Optical density was measured at 532 nm by Cakmak et al. [27].

### Hydrogen peroxide content (HPOC)

Fresh foliar material (0.5 gm) was chopped in 5 ml trichloro acetic acid (TCA) and centrifuged for 15 minutes; 0.5 ml supernatant was taken, and 0.5 ml phosphate buffer and 1 ml potassium iodide (KI) reagent was added to it. Optical density was recorded at 390 nm by following the methodology of [28].

### Endogenous tocopherol content (ETPC)

Methodology of Backer et al. [29] was followed to measure the levels of endogenous tocopherol content in leaf tissues. Leaf material (0.1 g) was grounded in 10 ml solution (petroleum ether and ethanol 2:1.6 v/v) followed by centrifugation for 15 minutes. 1 ml supernatant was taken and mixed with 0.2 ml (2%) 2-dipyridyl in ethanol (v/v). The mixture was poured into cuvete and placed in spectrophotometer. Optical density was noted at 520 nm.

### Ascorbate peroxidase activity (APX)

Ascorbate peroxidase (APX) levels were evaluated by pursuing the method of [30]. Fresh leaf material (0.5 gm) was grounded in 5 ml phosphate buffer (pH 7.0). The samples were centrifuged for 15 minutes and 0.2 ml supernatant was collected. 0.1mM hydrogen peroxide, 0.6 mM ascorbic acid and 0.1 mM ethylenediamine tetraacetic acid (EDTA) was added to supernatant and optical density was recorded at 290 nm.

### Catalase activity (CAT)

Catalase activity was measured by following the protocol of [31]. Fresh foliar material (0.5gm) was homogenized in 5 ml buffer solution (pH 7.0). Mixture was centrifuged at 3000 rpm for 15 minutes. 0.1 ml supernatant was taken 1.9 ml phosphate buffer (50 Mm) and 0.1 ml H_2_O_2_ (5.9 mM) was added. Optical density was noted at 240 nm for 3 minutes.

### Superoxide dismutase activity (SOD)

Foliar material (0.5 gm) was grounded in 5 ml phosphate buffer. Mixture was centrifuged for 15 minutes. After centrifugation 5 ml methionine, 150 µl riboflavin, 24 µl nitro-blue-tetrazolium (NBT) was added to 0.1 ml supernatant. Optical density was recorded at 560 nm by following the method of [32].

### Peroxidase activity (POD)

Peroxidase (POD) activity was determined by following the methodology of [31]. Leaf material (0.5 gm) was chopped in 2 ml morpholino ethane sulphonic acid (MES). The samples were centrifuged and 0.1 ml supernatant was collected. 1.3 ml MES, 0.1 ml phenyl diamine and 1 ml hydrogen peroxide (30%) were added. OD was noted at 470 nm for 3 minutes via spectrophotometer.

### Statistical analysis

Statistical analysis were based on three factors; comprised of lentil cultivar (Punjab-2009), drought stress of 20 and 25 days and alpha tocopherol levels 100, 200 and 300 mg/L. Experiment was conducted in randomized complete block design (RCBD) with three replicates for each group and 15 seeds per replicate. SPSS Statistic 25 was used for determining mean, standard error, analysis of variance (ANOVA) and correlation.

## Results

### Growth responses under induced drought stress and tocopherol levels

Statistical analysis revealed a significant increase at (*P*≤0.05) in AGR, CGR RGR and LAI with exogenously applied α-tocopherol 200 mg/L in comparison with control group and rest of the treatments under 20-days of induced drought stress. On contrary, these parameters were affected negatively under 20-days of induced drought stress with no α-tocopherol treatment (Figure 1-a, b, c, d). NAR and CVG showed improvement at (*P*≤0.05) in control group and group with 100 mg/L α-tocopherol treatment (Figure 2-a, b). Furthermore, RSR showed significant improvement at (*P*≤0.05) with α-tocopherol 200 mg/L and affected adversely on exposure to induced drought stress with no α-tocopherol application (Figure 2-c). In contrast with other treatments RWC was recorded maximum in the control group only (Figure 2-d). In comparison with the rest of the treatments and control group LAR showed significant results at (*P*≤0.05) with 200 mg/L α-tocopherol. On contrary, LAR was adversely affected under induced water stress condition with no α-tocopherol treatment. SVI showed positive response at (*P*≤0.05) in control group only (Figure 3-a, b). Same growth parameters were studied under 25-days of induced drought stress sprayed with the same levels of α-tocopherol. Results showed significant improvement at (*P*≤0.05) in AGR, LAI, and LAR with the application of 200 mg/L α-tocopherol only. On the other hand, AGR, CGR, LAI, LAR and NAR showed negative responses under induced drought stress of 25-days with no α-tocopherol application. NAR and CVG values were calculated highest for control group and 100 mg/L α-tocopherol treatment in contrast with the rest of the treatments.

**Figure 1.**
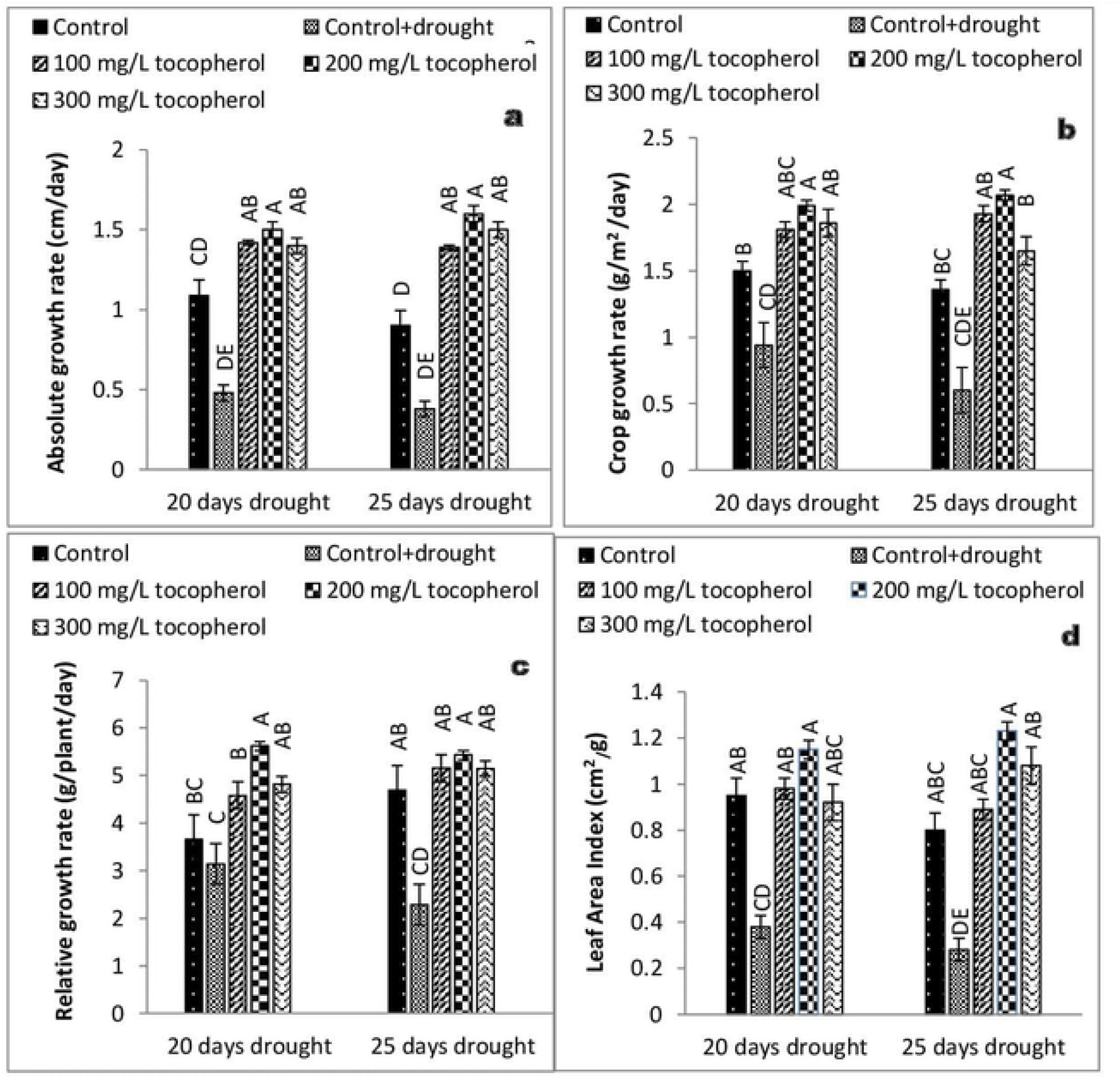
Effect of varying levels of exogenously applied α-tocopherol on absolute growth rate (**a**) crop growth rate (**b**) relative growth rate (**c**) leaf area index (**d**) of lentil (*Lens culinaris* Medik.) grown under varying drought stress condition (Mean ± standard error) letters (A–E) indicating least significance difference among the mean values at *p*≤0.05.

**Figure 2.**
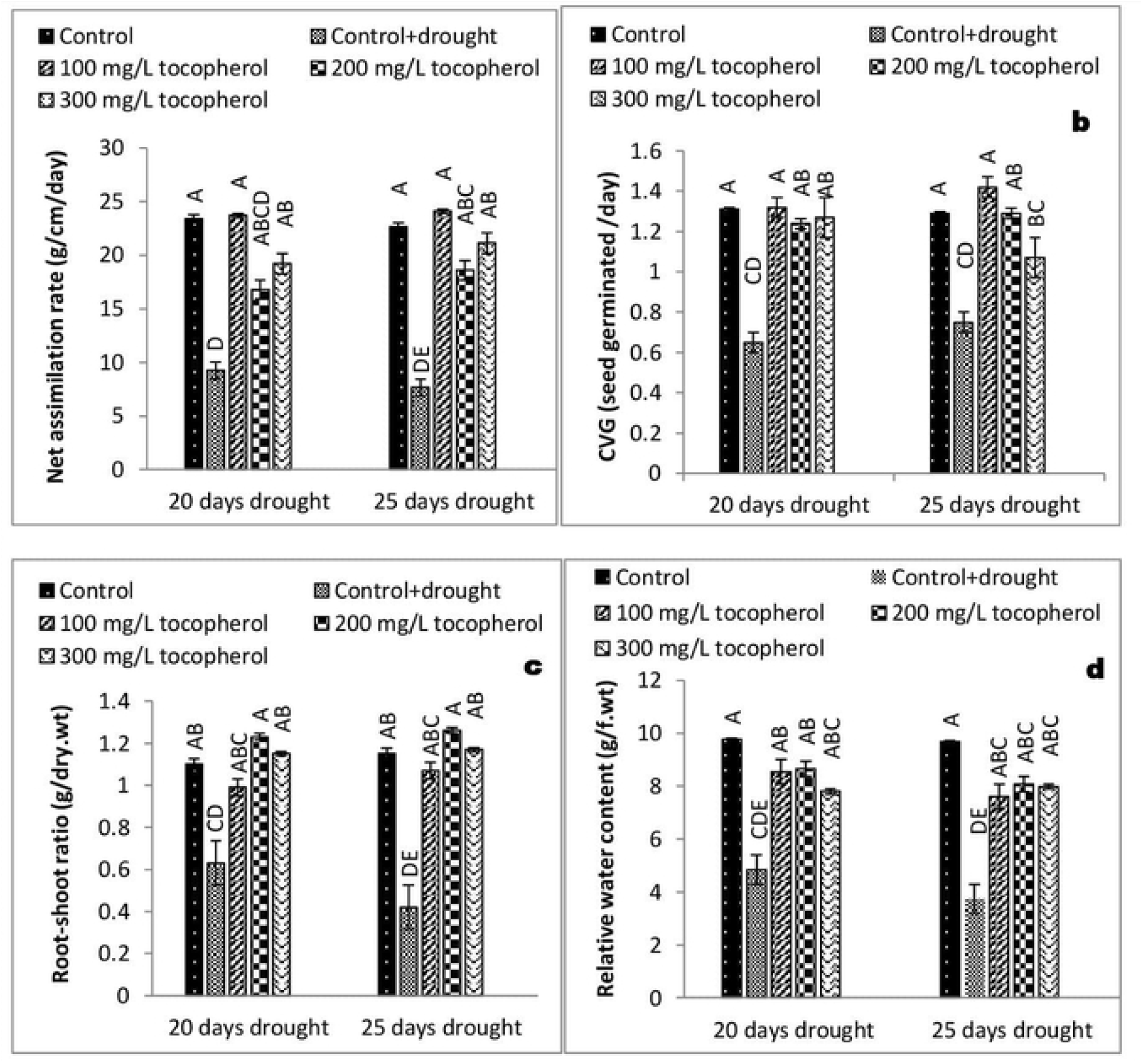
Effect of varying levels of exogenously applied α-tocopherol on net assimilation rate (**a**) coefficient of velocity of germination (**b**) root-shoot ratio (**c**) relative water content (**d**) of lentil (*Lens culinaris* Medik.) grown under varying drought stress condition (Mean ± standard error) letters (A–E) indicating least significance difference among the mean values at *p*≤0.05.

**Figure 3.**
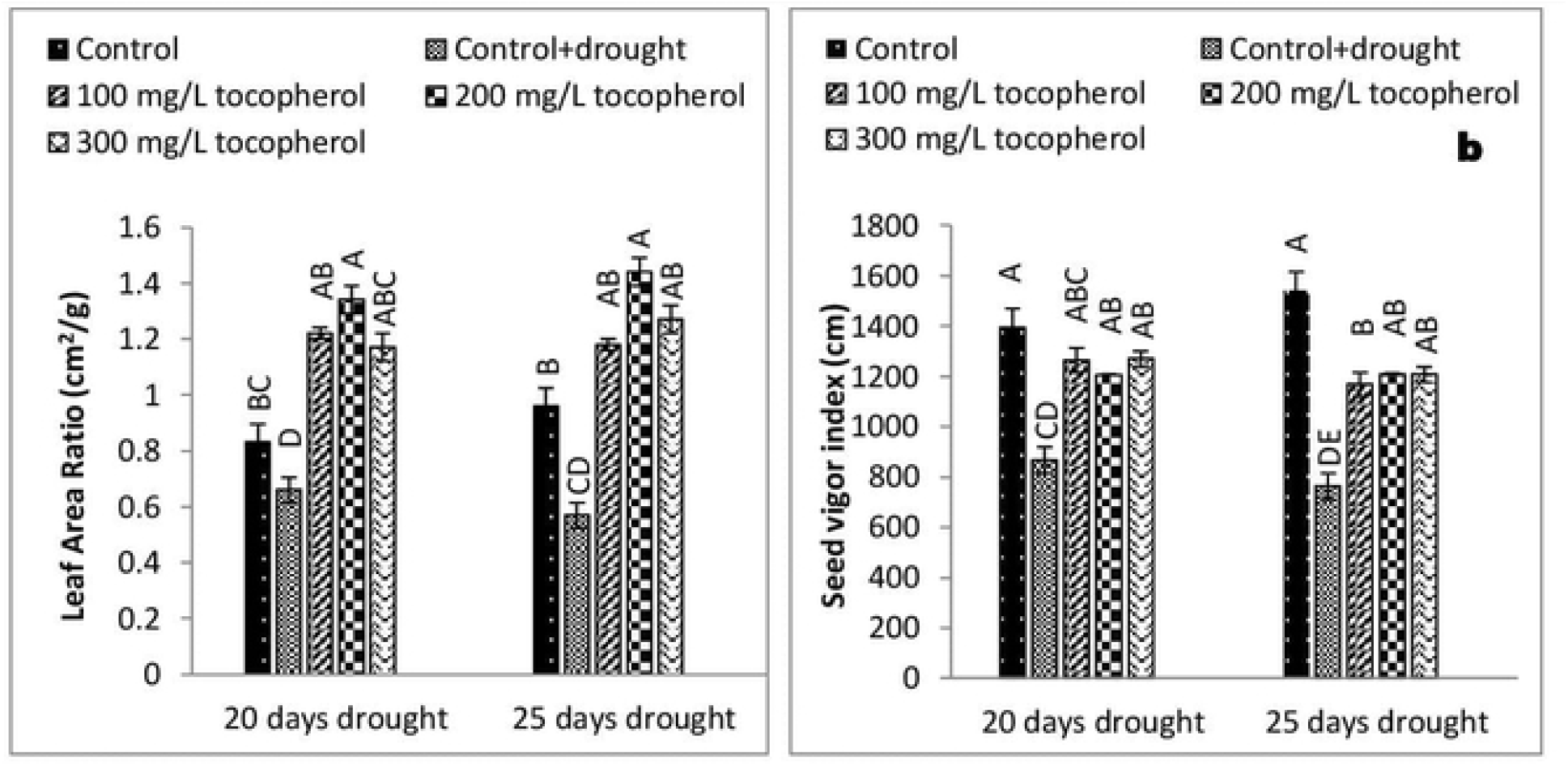
Effect of varying levels of exogenously applied α-tocopherol on leaf area ratio (a) seed vigor index (b) of lentil (*Lens culinaris* Medik.) grown under varying drought stress condition (Mean ± standard error) letters (A–E) indicating least significance difference among the mean values at p≤0.05.

RGR and RSR both showed significant enhancement at (*P*≤0.05) with the application of 200 mg/L α-tocopherol, both the parameters affected negatively on exposure to 25-days of induced drought stress with no α-tocopherol treatments. Subsequently, in comparison with the rest of treatments RWC and SVI values were highest only in the control group.

### Determination of photosynthetic pigments (Chlorophyll a, b & carotenoids)

Varying levels of induced drought stress markedly (*P*≤0.05) reduced chlorophyll “a”content, application of α-tocopherol enhanced (*P*≤0.01) chlorophyll “a” content in both the induced drought stress levels. In contrast with control group and rest of the treatments group sprayed with 200 mg/L tocopherol showed maximum chlorophyll “a”content (Table 1; Figure 4-a). Drought stress condition significantly (P≤0.01) decreased chlorophyll “b” content, α tocopherol ameliorated (*P*≤0.05) the levels of chlorophyll “b” content: among all the treatments and control group application of α-tocopherol 200 mg/L showed better response in improving the levels of chlorophyll “b” content (Table 1; Figure 4-b).

**Table 1.**
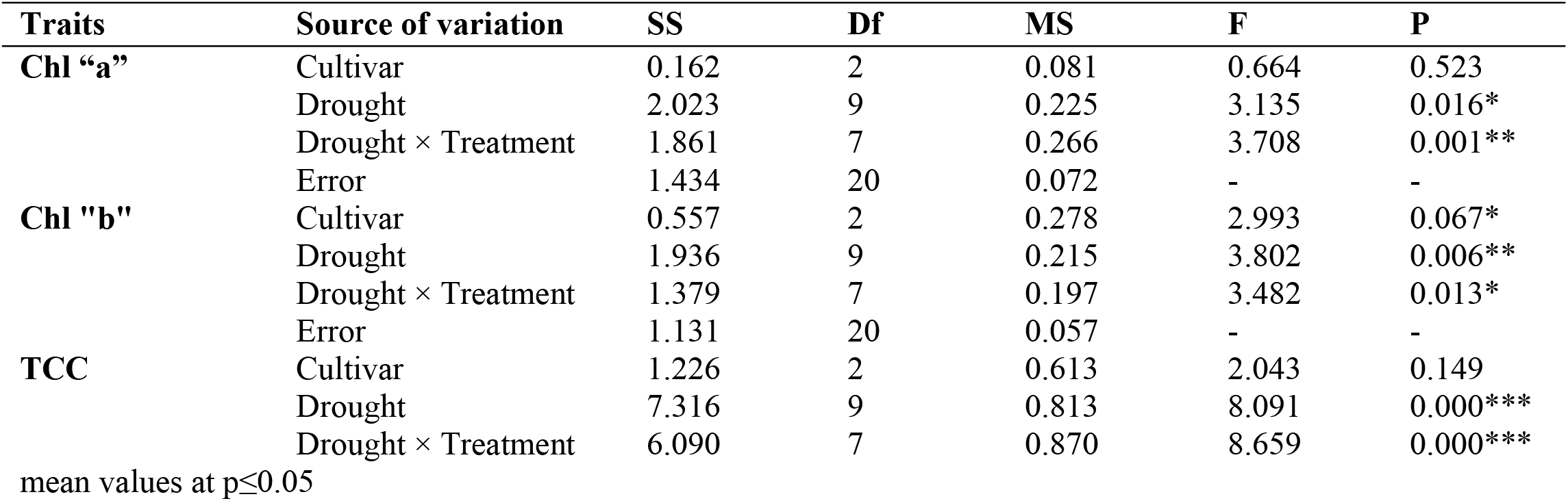
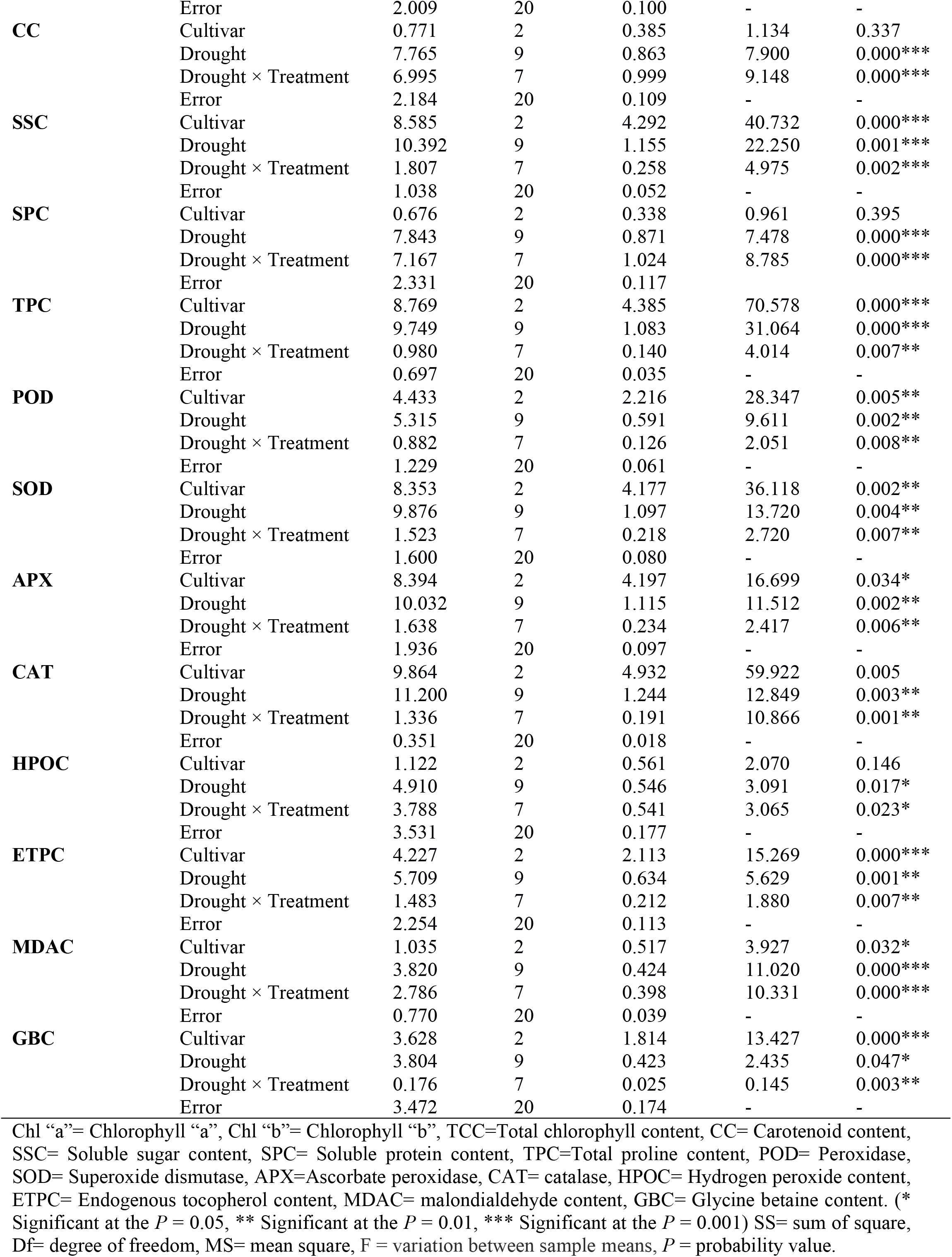
Analysis of variance of physio-biochemical attributes of lentil cultivar to varying levels of α-tocopherol under induced drought stress.

**Figure 4.**
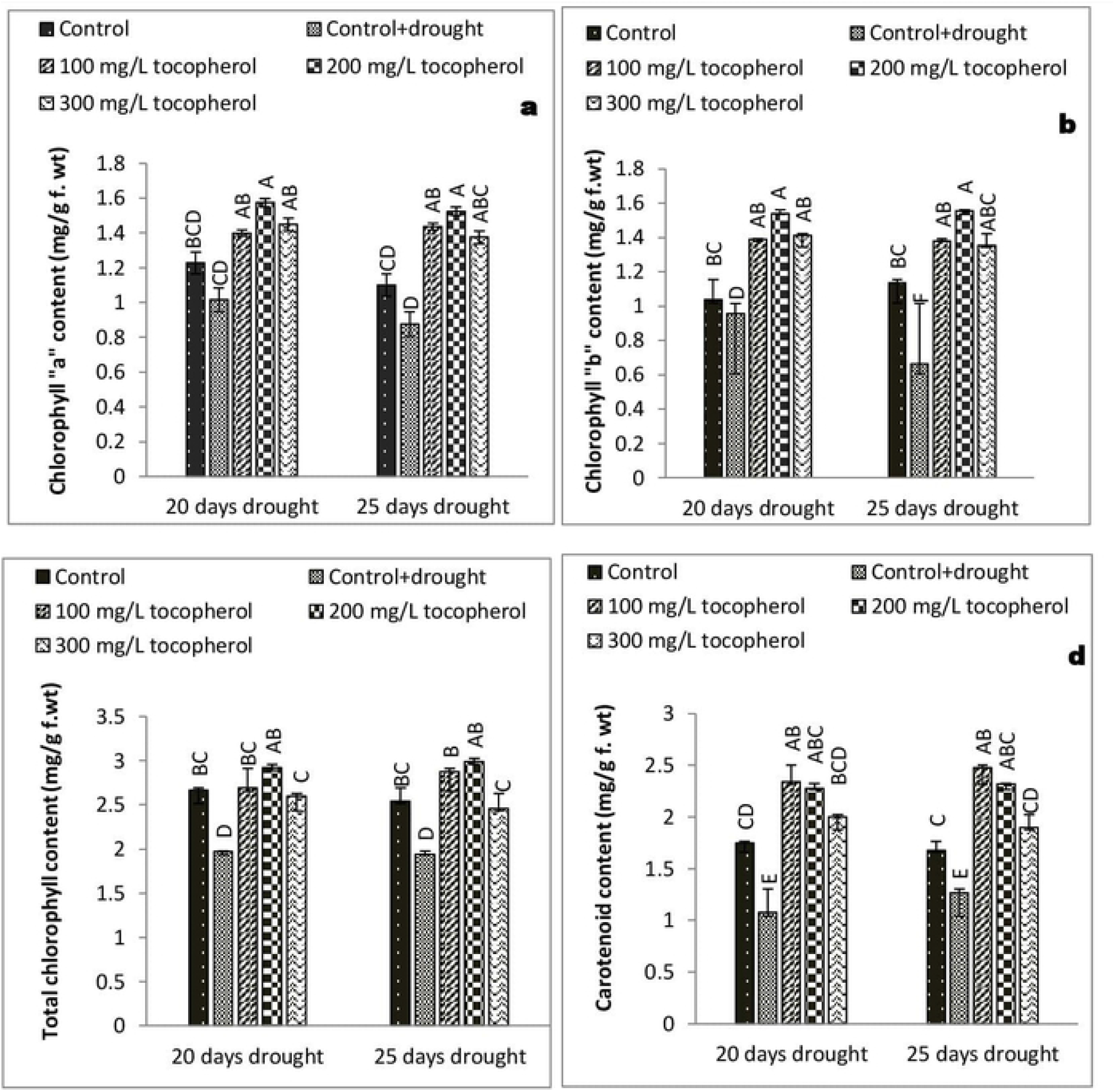
Effect of varying levels of exogenously applied α-tocopherol on chlorophyll “a” content (a) chlorophyll “b” content (b) total chlorophyll content (c) carotenoid content (d) of lentil (Lens culinaris Medik.) grown under varying drought stress condition (Mean ± standard error) letters (A–F) indicating least significance difference among the mean values at p≤0.05.

Induced drought stress regimes considerably (*P*≤0.001) reduced the levels of total chlorophyll content, among all the treatments and control group α-tocopherol 200 mg/L boosted (*P*≤0.001) the levels of total chlorophyll content in both (20 and 25 days) of induced drought stress levels (Table 1; Figure 4-c). Drought stress condition reduced (*P*≤0.001) the levels of carotenoid content to a great extent; as compared to other treatments and control group 100 mg/L of α-tocopherol application showed significant (*P*≤0.001) response in increasing the levels of carotenoid content (Table 1; Figure 4-d).

### Changes in concentration of soluble sugar content (SSC)

Drought stress condition had a significant influence on the concentration of soluble sugar content, limited water regimes raised (*P*≤0.001) the levels of soluble sugar content in both the drought levels. Exogenously applied α-tocopherol further raised (*P*≤0.001) the levels of soluble sugar content; in comparison with control and other applied treatments α-tocopherol 200 mg/L proved more effective in ameliorating soluble sugar content (Table 1; Figure 5-a).

**Figure 5.**
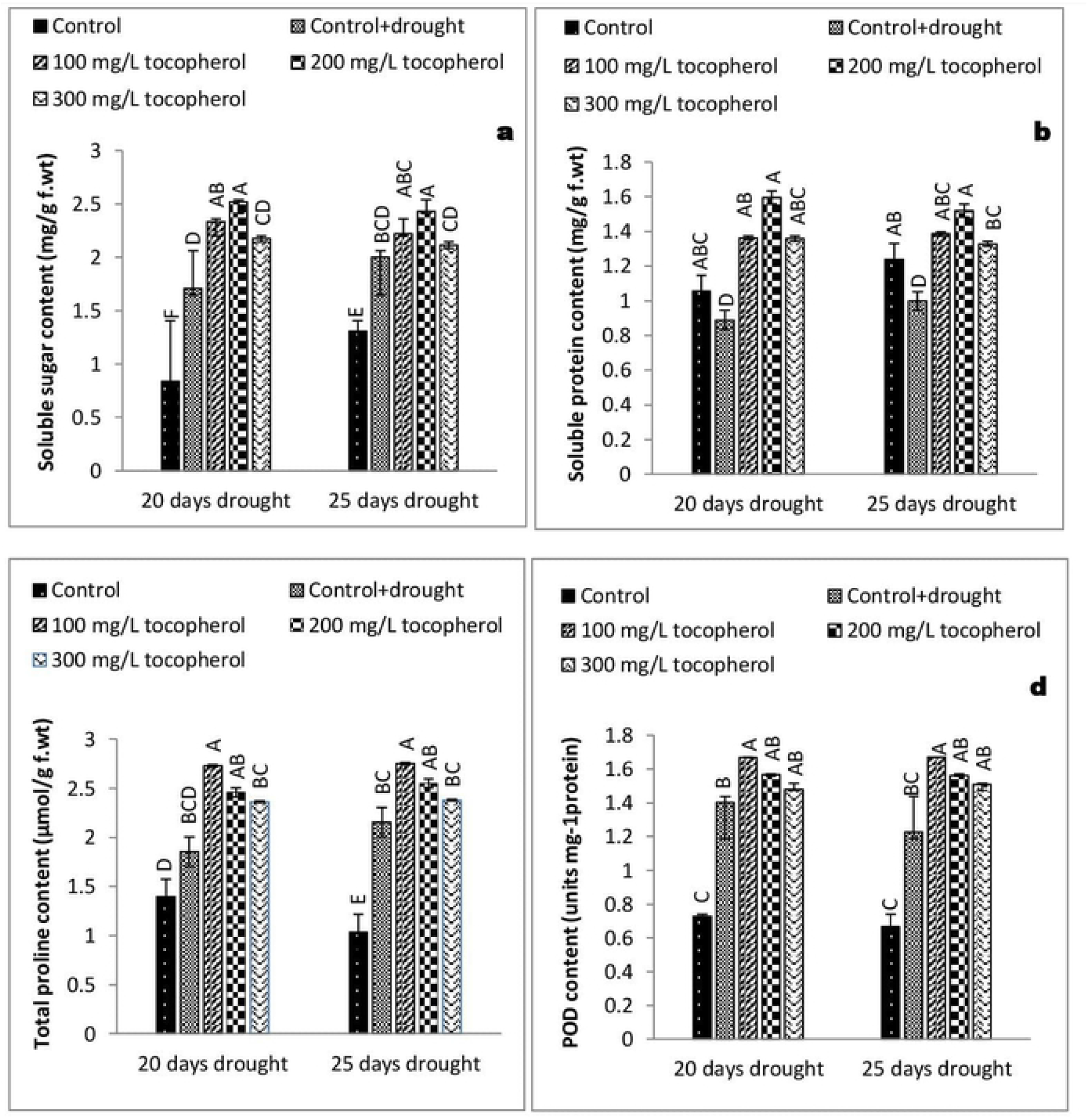
Effect of varying levels of exogenously applied α-tocopherol on soluble sugar content (a) soluble protein content (b) total proline content (c) peroxidase content (d) of lentil (Lens culinaris Medik.) grown under varying drought stress condition (Mean ± standard error) letters (A–F) indicating least significance.

### Effects on soluble protein content (SPC)

Drought stress inflicted a profound effect on soluble protein content as it markedly reduced (*P*≤0.001) the levels of soluble protein content. Among all the treatments α-tocopherol 200 mg/L was distinctly significant in raising the levels of soluble protein content (Table 1; Figure 5-b).

### Fluctuations in the levels of total proline content (TPC) and Glycine betaine content (GBC)

Drought stress condition condition increased total soluble proline content to a considerable level (*P*≤0.001). Application of α-tocopherol further enhanced (*P*≤0.01) soluble proline content under varying drought stress regimes, among different α-tocopherol levels 100 mg/L showed better results (Table 1; Figure 5-c).

### Responses of antioxidant enzymes activities (POD, SOD, APX & CAT)

On exposure to induced drought stressed condition a significant increase was noted in the activities of POD (*P*≤0.01), (Table 1; Figure 5-d). SOD (P≤0.01) APX (*P*≤0.01) and CAT (*P*≤0.01). Foliar application of α-tocopherol further significantly (*P*≤0.01) enhanced the activities of these enzymes. In case of peroxidase and superoxide dismutase 100 mg/L α-tocopherol treatment proved more effective. While, in case of ascorbate peroxidase and catalase 200 mg/L α-tocopherol responded better as compared to the rest of the treatments and control group (Table 1; Figure 6-a, b, c).

**Figure 6.**
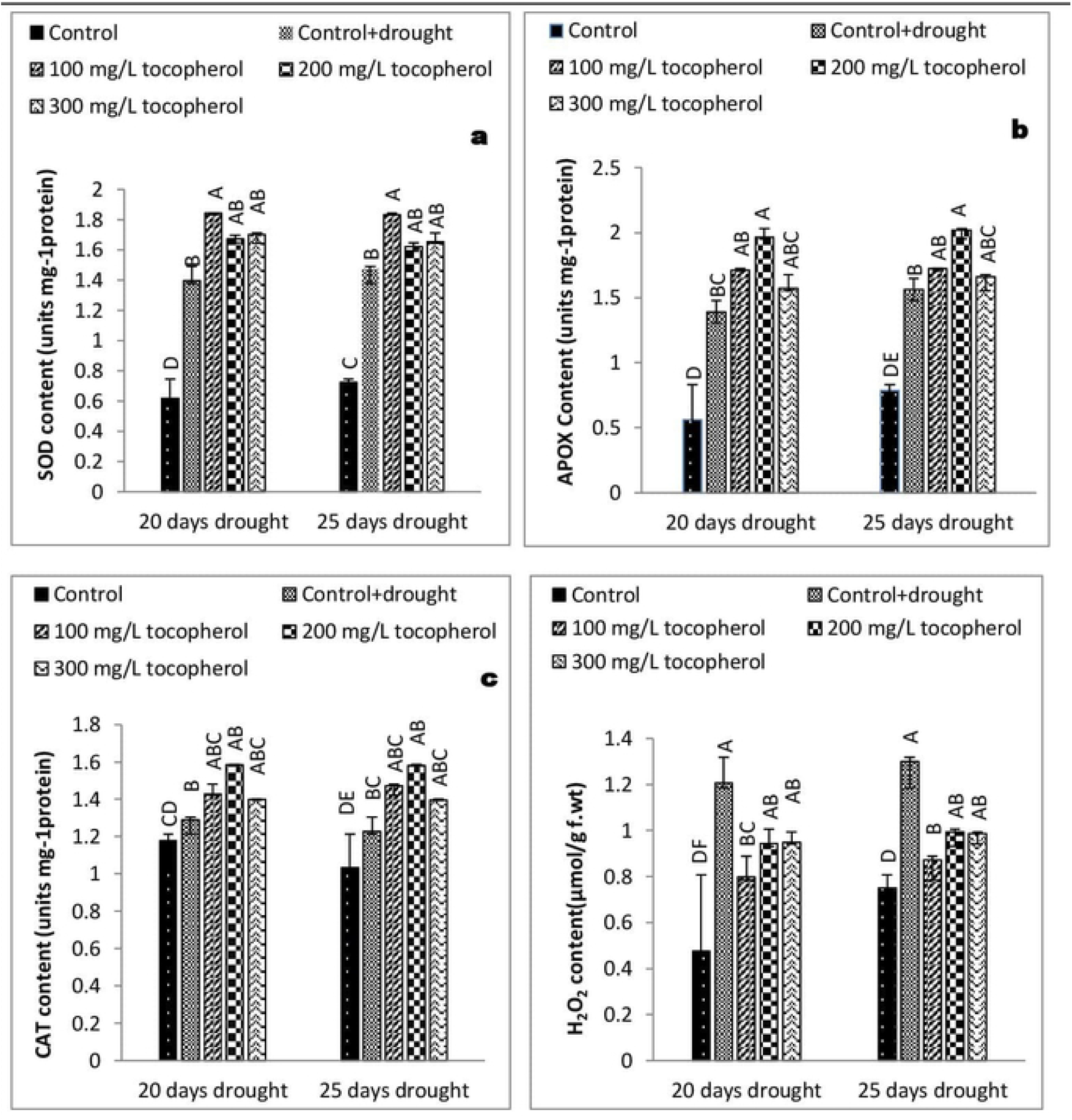
Effect of varying levels of exogenously applied α-tocopherol on superoxide dismutase content (a) ascorbate peroxidase content (b) catalase content (c) hydrogen peroxide content (d) of lentil (Lens culinaris Medik.) grown under varying drought stress condition (Mean ± standard error) letters (A–E) indicating least significance difference among the mean values at p≤0.05.

### Changes in the concentration of hydrogen peroxide (H_2_O_2_)

Drought stress condition caused a marked (*P*≤0.05) increase in hydrogen peroxide concentration, in comparison with control group and rest of the applied treatments α-tocopherol 200 and 300 mg/L showed a significant (*P*≤0.05) response in alleviating the levels of hydrogen peroxide content under limited water condition (Table 1; Figure 6-d).

### Levels of endogenous tocopherol content (ETPC)

Drought stress condition resulted in the accumulation of endogenous/membrane bounded tocopherol content to a level significant at (*P*≤0.01). Among all the foliar applied α-tocopherol treatments 100 mg/L tocopherol increased the levels of endogenous tocopherol content to a significant (*P*≤0.01) level (Table 1; Figure 7-a).

**Figure 7.**
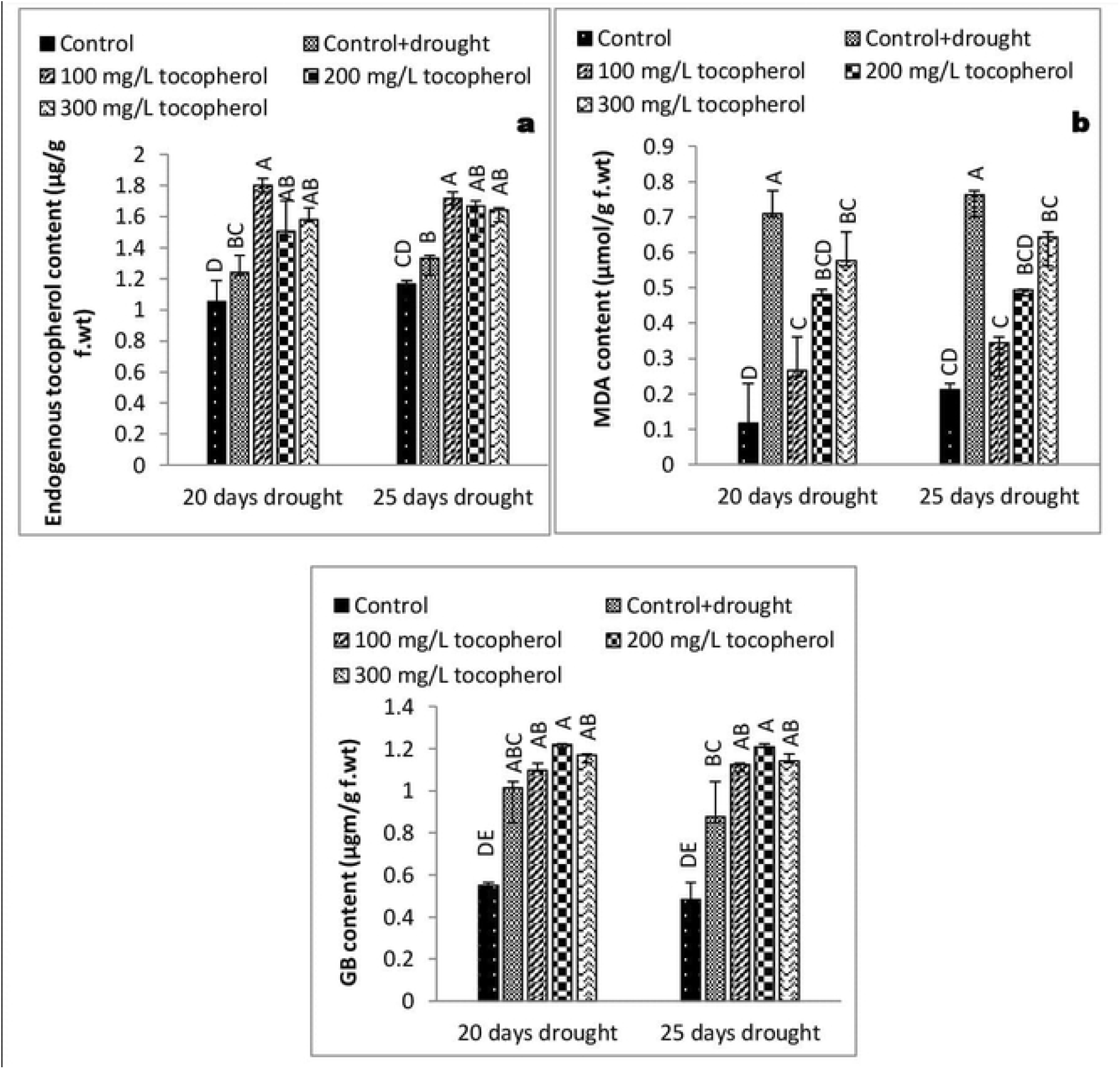
Effect of varying levels of exogenously applied α-tocopherol on endogenous tocopherol content (a) malondialdehyde content (b) glycine betaine content (c) of lentil (Lens culinaris Medik.) grown under varying drought stress condition (Mean ± standard error) letters (A–E) indicating least significance difference among the the mean values at p≤0.05.

### Changes in the concentration of Malondialdehyde (MDAC) content

Water deficit stress significantly (P≤0.001) raised the concentration of malondialdehyde content. On contrary, α-tocopherol application reduced the concentration of malondialdehyde to a considerable level (*P*≤0.001). However, in both the drought levels α-tocopherol 100 mg/L showed better response in decreasing the concentrations of malondialdehyde (Table 1; Figure 7-b).

### Fluctuations in the levels of total Glycine betaine content (GBC)

Water deficiency triggered a marked (*P*≤0.05) increase in glycine betaine content; foliar applied α-tocopherol further improved the concentration of glycine betaine content; α-tocopherol 200 mg/L showed better response in increasing the concentrations of glycine betaine content (Table 1; Figure 7-c).

### Correlation between induced drought stress, α-tocopherol and physio-biochemical attributes

A positively significant correlation was noted between varying drought stressed condition and physio-biochemical attributes of lentil including malondialdehyde, hydrogen peroxide, glycine betaine, soluble sugar, total proline, endogenous tocopherol contents and activities of antioxidant enzymes including peroxidase, superoxide dismutase, catalase and ascorbate peroxidase. Whereas; chlorophyll a, b, total chlorophyll content, carotenoid content and total soluble protein content correlated negatively towards both 20 and 25 days of induced drought stressed regimes. Although, a positive and significant correlation was observed between varying levels of exogenously applied α-tocopherol and chlorophyll a, b, total chlorophyll content, carotenoid content, soluble sugar content, soluble protein content, total proline content, endogenous tocopherol content, glycine betaine content and activities of antioxidant enzymes (APX, CAT, SOD and POD). However, a significantly negative correlation was observed between α-tocopherol levels and concentrations of hydrogen peroxide and Malondialdehyde content (Table 2).

**Table 2.** Correlation between induced drought stress, α-tocopherol and physio-biochemical attributes of lentil cultivar.

| **Correlation analysis between drought stress and physio-biochemical attributes** |  |  |  |  |  |  |  |  |  |  |  |  |  |  |
| --- | --- | --- | --- | --- | --- | --- | --- | --- | --- | --- | --- | --- | --- | --- |
| Chl "a" | Chl "b" | TCC | CC | SSC | SPC | TPC | POD | SOD | APX | CAT | HPOC | ETPC | MDAC | GBC |
| -0.421* | -0.562* | -0.601* | -0.654* | 0.862* | -0.854* | 0.907* | 0.799* | 0.885* | 0.854* | 0.833* | 0.968* | 0.777* | 0.961* | 0.882* |
| **Correlation analysis between tocopherol and physio-biochemical attributes** |  |  |  |  |  |  |  |  |  |  |  |  |  |  |
| Chl "a" | Chl "b" | TCC | CC | SSC | SPC | TPC | POD | SOD | APX | CAT | HPOC | TPC | MDAC | GBC |
| 0.893* | 0.884* | 0.899* | 0.903* | 0.911* | 0.941* | 0.993* | 0.887* | 0.918* | 0.889* | 0.924 | -0.551* | 0.889* | -0.647* | 0.919* |

Chl "a"= Chlorophyll "a", Chl "b"= Chlorophyll "b", TCC=Total chlorophyll content, CC= Carotenoid content, SSC= Soluble sugar content, SPC= Soluble protein content, TPC=Total proline content, POD= Peroxidase, SOD= Superoxide dismutase, APX=Ascorbate peroxidase, CAT= catalase, HPOC= Hydrogen peroxide content, ETPC= Endogenous tocopherol content, MDAC= malondialdehyde content, GBC= Glycine betaine content. (\* Significant at the $P = 0.05$ , \*\* Significant at the $P = 0.01$ , \*\*\* Significant at the $P = 0.001$ )

## Discussion

### Growth responses

Oxidative stress and higher production of ROS have been considered the major causes that adversely affect plant growth [33]. Reduction in plant growth parameters such as LAI and LAR have been reported to be mainly caused by stomatal closure during drought stress condition, that eventually leads to leaf senescence [34]. On exposure to drought stress condition plants tend to close their stomata which slow down the water loss from aerial parts of the plants; as a result the carbon dioxide (CO^2^) absorption is reduced and consequently net photosynthesis [35]. Moreover, reduction in growth attributes such as CVG, CGR, AGR, RGR, RSR, SVI and NAR are closely linked with suppression of cell elongation and cell growth under drought stress regimes [36]. In the present study induced water deficit stress impacted negatively on lentil growth and physio-biochemical mechanisms; on contrary foliar applied α-tocopherol curbed the negative consequences of induced drought stress by regulating key metabolic activities and boosted the growth parameters including AGR, CGR, LAI, LAR, NAR and CVG (Figures. 1, 2, 3). Induced water deficit stress caused a considerable reduction in AGR, CGR, LAI, LAR and NAR of lentil. Likewise, Teimouri et al, [37], Riccardi et al, [38] also observed reduction in these growth parameters under induced drought stress conditions. It is believed that this reduction is due to stomatal closure, persistent exposure to limited water regimes ultimately leading to shrinkage of leaves by Teimouri et al, [37] and Zaheer et al. [39]. Similar variations in growth parameters were recorded by other workers [40, 41]. Results from statistical analysis in (Figures. 1, 2, 3) indicated that RSR, RGR, RWC and SVI were improved in the control group and affected negatively in groups subjected to varying drought stress levels with no α-tocophrol treatments; thus, confirmed the same investigation by [42].

### Physiological and biochemical responses

Under drought stress conditions an increase in chlorophyll degrading enzymes activity prompts the destruction of chlorophyll pigments [43]. Drought stress causes chlorophyll pigments degeneration and hampering the process of photosynthesis [44, 45]. Chloroplast is the most sensitive organelle to drought stress exposure, and it has been proved that drought stress damages photosynthetic pigments in various crops [46, 47].

It is suggested that the degeneration of chlorophyll is associated with the production of ROS, which lowers down the rate of photosynthesis and increases cellular respiration [48, 49]. In the present study photosynthetic pigments (chlorophyll a, b and carotenoid content) affected negatively on exposure to induced drought stress. In contrast foliar applied α-tocopherol significantly enhanced the levels of photosynthetic pigments (Fig. 4a-d) Our results were in accordance with the previous research work carried out by [9, 50, 51].

In response to abiotic stress, plants perceive a disturbance in their physiological activities and respond abruptly by accumulating a variety of osmolytes mainly including glycine betaine and proline; osmolytes provide conducive environment for various metabolic activities and protecting plants from the damages caused by oxidative stress described by [52]. Compounds like proline, MDA and H_2_O_2_ are generally being used as stress markers, proline is well known for its osmoprotective role, in many plants an increase concentration of proline during drought stress condition indicated to be correlated with drought stress tolerance [53]. It’s also evident that proline has the potential to directly act as ROS scavenger and regulator of cellular redox status [54]. The studied lentil cultivar revealed a marked rise in soluble sugar content (Fig. 5-a), total proline content (Fig. 5-c), and glycine betaine content (Fig. 7-c) by experiencing water deficit stress condition. Similarly, foliar applied α-tocopherol further increased the concentrations of these osmolytes; our results were parallel with the investigation made by [55] in chinese rye grass (*Leymus chinensis*), a high level of proline was observed in drought stressed seedlings grown from seeds primed with α-tocopherol. The same results were recorded by [56] in soybean (*Glycine max*) and in faba bean (*Vicia faba)* by [57].

Water limited condition cause protein and lipid degeneration and affect plant growth and other vital activities [58]. In case of lentil, it was found that low moisture content in the soil alleviated the soluble protein content to a great extent. Application of α-tocopherol 200 mg/L was distinctly significant in raising the levels of soluble protein content (Fig. 5-b). Sadak and Dawood, [51] concluded that flax (*Linum usitatissimum*) plants sprayed with α-tocopherol under salinity stress caused a marked increase in soluble protein content. Similarly,

[57] suggested that application of 100 mg/L of α-tocopherol in faba bean (*Vicia faba*) plants proved significant in preservation of soluble protein content.

The main role of *a*-tocopherol is the removal of lipid peroxyl radical prior of its attack to target lipid substrate synthesizing *a*-tocopheroxyloxyl radicals [59]. Under abiotic stress condition a-tocopherol deactivates ^1^O_2_ in chloroplast, according to an estimate a single molecule of a-tocopherol can deactivate 120 molecules of ^1^O_2_ [60]. Membrane bounded tocopherol levels were observed maximum in lentil plants exposed to drought stress regimes. Application of α-tocopherol 100 mg/L proved more significant in raising the levels of membrane bounded tocopherol content (Fig. 7-a). Our findings in the case of lentil were supported by the investigations made by Ali et al. [61] who reported high levels of endogenous tocopherol contents in maize plants cultivated under induced water stress condition. Zhang et al. [62] confirmed same results in canna (*Canna edulis*) cultivars under induced drought stress condition.

Malondialdehyde (MDA) is the product of membrane degradation; under abiotic stress condition a rise in MDA concentration marks the disintegration of biological membrane [63]. Accumulation of MDA has been considered an indication of lipid peroxidation in various plants under stress condition [64, 65]. In plant tissues peroxidation of free fatty acid could occur both in non-enzymatic and enzymatic ways, producing a number of breakdown products which mainly include alcohol, aldehydes and their esters and this process is considered to be mainly involved in oxidative damage to cellular membranes and other biomolecules [66]. Physiological analysis of lentil cultivar revealed an increase in MDA content under drought stress condition. Although, exogenously applied α-tocopherol significantly decreased the levels of MDA content (Fig. 7-b). The same results were obtained by [67] in case of geranium (*Pelargonium graveolens*) treated with 100 mg/L α-tocopherol.

Hydrogen peroxide (H_2_O_2_) being a prominent reactive oxygen species (ROS) in plants results in cell oxidation, disturbs vital metabolic processes and interrupts membrane stability under stress condition [68]. Plants naturally produce ROS, mainly including H_2_O_2_ superoxides, there is a delicate balance between ROS production and it’s scavenging, under drought stress condition this balance is disturbed as plant tend to close their stomata which also limits CO2 fixation [69]. In present study, a significant increase was observed in H_2_O_2_ content under drought stress condition. Application of α-tocopherol decreased the concentration of H_2_O_2_ to a considerable level (Fig. 6-d). Parallel to our results Kostopoulou et al. [70] recorded that application of resveratrol and α-tocopherol decreased the levels of H_2_O_2_ in citrus plants subjected to salinity stress. The same results were reported by [71] who found that 100 mg/L α-tocopherol proved better in lowering the levels of H_2_O_2_ in wheat plants exposed to salinity stress.

Plants counteract the oxidative stress generated by ROS in a coordinated way both enzymatically and non-enzymatically [72, 73]. SOD acts as first line of defence as it converts the superoxide radicals to H_2_O_2_ [74, 45]. Likewise, Hojati et al. [75] in their demonstration on *Carthamus tinctorius* cultivars subjected to drought stress, concluded that enzymatic as well as non-enzymatic antioxidants were involved in the removal of ROS, SOD is needed to scavenge superoxide radical [76], while scavenging H_2_O_2_, requires POD, CAT and APX [77]. Natural self defence systems are well developed in plants; on exposure to stress condition these defence systems are activated both enzymatically (Ascorbate peroxidase, catalase, superoxide dismutase, peroxidase) and non-enzymatically (secondary metabolites) scavenging the reactive oxygen species (ROS) formed as result of stress condition [73]. Though, activities and response of these antioxidants varies plant to plant [75]. In the present research study, water stress condition caused a prominent increase in the activities of antioxidant enzymes POD

(Fig. 5d), SOD, APX and CAT (Fig. 6-a, b, c). However, foliar applied α-tocopherol further enhanced the activities of these enzymes. Our results were in agreement with findings made by [55] who observed a marked increase in SOD and POD activities in chinese rye grass (*Leymus chinensis*) subjected to induced water stress condition. Similarly, Rady et al, [78] reported an increase in the activities of POD and CAT in sunflower plants under salt stress. Furthermore, [57] reported that, application of α-tocopherol 200 mg/L enhanced the performance of antioxidant enzymes to a great extent in drought stress plants of faba beans (*Vicia faba*).

## Conclusions

From the present study it was confirmed that exogenously applied α-tocopherol under induced water deficit stress regimes ameliorated drought stress tolerance potential of lentil cultivar to a great extent; by enhancing growth, physiological and biochemical attributes. Furthermore, application of α-tocopherol 200 mg/L followed by 100 mg/L showed better response in mitigating the damaging effects of induced drought stress. Importantly, the present scenario of alarmingly increasing changes in climatic conditions appeals the researchers that still there is a dire need to further investigate various growth and physiological responses of lentil cultivars to different types of biotic and abiotic stresses.

## Acknowledgements

We are highly acknowledged to the Department of Botany, University of Peshawar for providing all facilities regarding this work.

## Data Availability

Data associated with the present paper can be obtained by contacting the corresponding authors.

## Authors’ contributions

Sami Ullah designed the experiments and supervised the experiments of the first author Wadood Shah who conducted the experiments and analysed all data. Sajjad Ali, Muhammad Idrees and Muhammad Nauman Khan conduct antioxidants and biochemical analysis, Kasif Ali^4^, Ajmal Khan did statistical analysis and help in manuscript writing, Muhammad Ali and Frhan Younas reviewed and checked the final version. All authors have read and agreed to its contents and also that the manuscript complies with the policy of the publication.

## Abbreviations

AGR: Absolute growth rate
APX: Ascorbate peroxidase
CAT: Catalase
GB: Glycine betaine
H_2_O_2_: Hydrogen peroxide
LAI: Leaf area index
LAR: Leaf area ratio
MDA: Malondialdehyde
POD: Peroxidase
ROS: Reactive oxygen species
RGR: Relative growth rate
RSR: Root shoot ratio
SOD: Superoxide dismutase
SPC: Soluble protein content
SSC: Soluble sugar content
TPC: Total proline content
CGR: Crop growth rate
NAR: net assimilation rate
CVG: coefficient of velocity of germination
RWC: relative water content
SVI: seed vigor index.

